# Chronic Psychosocial Stress and Experimental Pubertal Delay Affect Socioemotional Behavior and Amygdala Functional Connectivity in Adolescent Female Rhesus Macaques

**DOI:** 10.1101/2020.09.18.304048

**Authors:** Melanie Pincus, Jodi S. Godfrey, Eric Feczko, Eric Earl, Oscar Miranda-Dominguez, Damien Fair, Mark E. Wilson, Mar M. Sanchez, Clare Kelly

## Abstract

In females, pubertal onset appears to signal the opening of a window of increased vulnerability to the effects of stress on neurobehavioral development. What is the impact of pubertal timing on this process? We assessed the effects of pubertal timing and stress on behavior and amygdala functional connectivity (FC) in adolescent female macaques, whose social hierarchy provides an ethologically valid model of chronic psychosocial stress. Monkeys experienced puberty spontaneously (*n*=34) or pubertal delay via Lupron treatment from age 16-33 months (*n*=36). We examined the effects of stress (continuous dimension spanning dominant/low-stress to subordinate/high-stress) and experimental pubertal delay (Lupron-treated vs. Control) on socioemotional behavior and FC at 43-46 months, after all animals had begun puberty. Regardless of treatment, subordinate monkeys were more submissive and less affiliative, and exhibited weaker FC between amygdala and dorsolateral prefrontal cortex and stronger FC between amygdala and temporal pole. Regardless of social rank, Lupron-treated monkeys were also more submissive, less affiliative, and explored less in a “Human Intruder” task but were less anxious than untreated monkeys; they exhibited stronger FC between amygdala and orbitofrontal cortex. No interactions between rank and Lupron treatment were observed. These data suggest that some of the effects of chronic subordination stress and delayed puberty overlap behaviorally, such that late-onset puberty-linked exposure to female hormones mimics chronic stress. In the brain, however, delayed puberty and subordination stress had separable effects, suggesting that the overlapping socioemotional outcomes may be mediated by distinct neuroplastic mechanisms. To gain further insights, additional longitudinal studies are required.

## 1. INTRODUCTION

Adolescence is a critical developmental period during which the brain is particularly sensitive to the effects of adverse experience such as chronic social stress (Dahl et al., 2018; Fuhrmann et al., 2015). Females appear to be at greater risk for deleterious psychological outcomes during adolescence than males (Thapar et al., 2012). One hypothesis is that sex-specific pubertal increases in gonadal hormones such as estradiol (E2) potentiate stress-induced plasticity within brain circuits supporting emotion regulation and cognition (McEwen and Morrison, 2013; Shansky et al., 2006). Such disruptions to typical neurodevelopment “take root,” later manifesting as psychopathology. What remains unclear is the role of timing in this process – the extent to which the timing of the intersection of stress with *developmentally mediated* plasticity (linked with chronological age) vs. *hormone-mediated* plasticity (linked with pubertal onset) confers risk for deleterious neurobehavioral outcomes.

The steep drop in the age of the onset of female puberty in developed nations (e.g., Herman-Giddens, 2017; 2006), combined with rising rates of depression and anxiety among young women (Twenge et al., 2019) suggests that early female puberty may confer increased vulnerability to stress-linked psychopathology, likely reflecting both biological and psychosocial mechanisms (Copeland et al., 2019). Other data suggest that chronological age is less important, however, and that the hormonal changes associated with puberty confer increased risk of mood dysregulation, regardless of timing (e.g., Lewis et al., 2018). Data linking contraceptive medication with increased depressive symptoms (Skovlund et al., 2016) and suicide risk (Skovlund et al., 2018), further suggest that elevated exposure to female gonadal hormones at any age increases vulnerability to mood dysregulation. A key challenge for studies examining these links is to separate chronological age from pubertal timing and stage, as these are inherently entwined and influenced by complex confounds, including body-mass index and socioeconomic status (Herman-Giddens, 2017). Further, studies on the role of stress in catalyzing negative behavioral outcomes are constrained to be correlational.

By permitting direct experimental manipulations of hormone exposure and stress, translational research with non-human animals overcomes these challenges. Such research is uncovering a picture of how gonadal hormones shape brain structure and function during adolescence. In rodents, sex differences in the number of neurons in ventromedial prefrontal cortex (vmPFC), and in vmPFC grey and white matter volume in adulthood (females < males) reflect differential rates of neuron and glial cell loss, pruning, and myelination during adolescence (e.g., Koss et al., 2015; Markham et al., 2007). When female rodents undergo prepubertal gonadectomy, these sexual dimorphisms do not emerge, suggesting that they are driven by puberty-related increases in ovarian hormones, including E2 (Koss et al., 2015). Evidence of the mechanisms through which pubertal E2 increases shape brain function and behavior is also emerging. For example, typical (puberty-linked) or early (pre-pubertal) but not late (post-pubertal) exposure to elevated E2 drives the maturation of inhibitory neurotransmission in mouse cingulate, thus altering the plasticity of cortical networks supporting cognitive control of behavior that are consistently implicated in psychopathology (Piekarski et al., 2017).

Although sex differences in the effects of stress on brain and behaviour in *adult* rodents are well documented (e.g., McEwen et al., 2016; Rincón-Cortés et al., 2019), murine research on how stress alters *neurodevelopment* has typically focused on males. Emerging studies on sex differences suggest that stress alters developmental plasticity in areas such as medial PFC, amygdala, and hippocampus in both sex-dependent and sex-independent ways within a specific time window triggered by puberty (e.g., Breach et al., 2019; Eiland et al., 2012). Increased female vulnerability to some of the effects of stress on socioemotional behaviors during adolescence (e.g., Bourke and Neigh, 2011; McCormick and Green, 2013; Weintraub et al., 2010) further suggests differential stress-related alteration of neurodevelopment in males and females. Together, these findings suggest that the normative pubertal rise in E2 may trigger a sex-determined window of increased female vulnerability to the effects of stress during adolescence (Hodes and Epperson, 2019; Naninck et al., 2011). Earlier puberty may thus widen or prematurely open the window of risk. A corollary hypothesis is that later onset of puberty may reduce risk, since the developing brain may become *less sensitive* to the effects of gonadal hormones with increasing age (Schulz and Sisk, 2016). Secular trends towards earlier puberty onset, rising rates of depression and anxiety among young women, and the growth in medical treatment of girls showing signs of precocious puberty (e.g., Eugster, 2019), highlight the importance of examining how the timing of exposure to puberty-linked increases in gonadal hormones interacts with stress in the developing brain.

Neuroimaging studies have begun to disentangle the effects of age, sex, and pubertal stage on brain structure and function in humans (e.g., Goddings et al., 2019; Herting et al., 2015; van Duijvenvoorde et al., 2019; Wierenga et al., 2018), as well as the impact of stress during development (e.g., Cohodes et al., 2020; Fareri et al., 2017; Tottenham and Galvan, 2016). Their findings map well onto the rodent work, demonstrating altered structure, function, and connectivity within and between regions such as amygdala, striatum, hippocampus, and PFC. Examining how pubertal hormones interact with stress remains a challenge for human studies, however, because it is not permissible to experimentally manipulate these. Studies with non-human primates offer a translational bridge from rodent work. In a recent example, Reding et al. (Reding et al., 2019) showed that, in ovariectomized adult female rhesus monkeys, treatment with E2 differentially modulated amygdala-medial PFC functional connectivity in chronically stressed (subordinate) relative to non-stressed (dominant) animals. This suggests that, even during adulthood, exposure to E2 interacts with chronic stress to alter macroscale brain circuits of relevance to human psychopathology.

Here, we examined a similar question in a neurodevelopmental context – adolescence – using the same macaque social subordination model. By experimentally delaying puberty onset in half our sample of adolescent female macaques using a pharmacological blocker, Lupron, we could dissociate chronological age from pubertal timing and compare the effects of species-typical (Control) vs. late (Lupron-treated) pubertal onset on stress-related brain and behavioral phenotypes. We expected that subordinate status would be associated with stress-related socioemotional impairments and altered functional connectivity in amygdala circuits, and contrasted the following opposing hypotheses regarding the impact of pubertal timing: (1) given evidence that elevated exposure to female gonadal hormones increases vulnerability to stress, regardless of age, subordination stress will have robust negative effects, regardless of chronological age at puberty onset (i.e., there will be no interaction between social status and pubertal delay), or, (2) since the brain may become less sensitive to the organizing effects of gonadal hormones with increasing age, experimentally delaying puberty will protect against the effects of subordination stress (i.e., social status and pubertal delay will interact, such that treated subordinates with late-onset puberty will exhibit fewer negative outcomes than subordinates with typical puberty). Our study provides the first experimental test of these two conflicting predictions concerning the role of chronological age vs. puberty-linked hormonal changes in determining the deleterious impact of stress on neurobehavioral development.

## 2. METHODS

### 2.1. Subjects

Subjects were 70 female rhesus macaques (*Macaca mulatta*) housed at the Yerkes National Primate Research Center (YNPRC) Field Station in Lawrenceville, GA, living in 4 large social groups made up of 2-3 adult males, 30-60 adult females, and their offspring. The animals were housed in outdoor enclosures with access to climate-controlled indoor facilities. They had *ad libitum* access to water and a standard low-fat, high-fiber diet (Purina Mills Int., Lab Diets, St. Louis, MO, USA) supplemented with fresh fruit and vegetables. The present analyses are part of a family of studies of this cohort examining the effects of social status on a range of neurobehavioral outcomes from the early juvenile, pre-pubertal-period through post menarche and first ovulation (e.g., Wilson et al., 2013). All proce**d**ures were approved by the Emory University Institutional Animal Care and Use Committee in accordance with the Animal Welfare Act and the U.S. Department of Health and Human Services “Guide for Care and Use of Laboratory Animals.”

### 2.2. Social rank

Socially housed female macaques organize into a linear social dominance hierarchy in which subordinate members are subject to constant unpredictable harassment, mimicking the uncontrollable and unpredictable nature of human psychosocial stress (Wilson, 2016). Subordinate group-living macaques experience continuous exposure to subordination stress from early in life and exhibit a range of stress-related neurobehavioral (Godfrey et al., 2016; e.g., Howell et al., 2014; Reding et al., 2019) and health outcomes (Sapolsky, 2005; e.g., Snyder-Mackler et al., 2016).

Our ethologically valid model operationalizes chronic psychosocial stress (PS) as subordinate social status, a continuous dimension spanning dominant social status (low-PS) to subordinate social status (e.g., Wilson et al., 2013). Each animal’s relative social rank was determined from the outcome of dyadic agonistic interactions between group mates, based on frequent 30-minute observations throughout the study period (Wilson et al., 2013). A more subordinate rank was assigned to an animal that unequivocally submitted to another animal. Each animal’s relative social rank is calculated as a number between 0 and 1 that represents her rank divided by the total number of animals in the social group, excluding animals <12 months old. Accordingly, dominant animals have *low relative ranks* (e.g., an animal ranked 1 out of 100 animals has a relative rank of 0.01) while subordinate animals have *high relative ranks* (e.g., an animal from the same group with a rank of 85 has a relative rank of 0.85). Relative ranks were consistent throughout the study for all but seven subjects who were removed from their natal group just before the 42-month data collection point because of intra-group aggression. For these subjects, their relative social rank at the time of removal was used, as it best reflects their life history and experience.

### 2.3. Lupron administration

Animals were randomly assigned to a Control group (*n*=36) that reached menarche spontaneously (30.88±0.64 months; Fig. 1) or to group that received depot Lupron treatment (Lupron-treated; *n*=34), resulting in delayed menarche (38.07±0.34 months). Depot Lupron is a sustained release gonadotropin-releasing hormone (GnRH) agonist that down-regulates GnRH receptors in the pituitary, suppressing developmental increases in gonadotropins and E2. The drug was administered monthly (0.25μg/kg/mo, i.m.) from 16.36±0.23 to 32.56±0.23 months of age, to experimentally delay menarche until a later chronological age. Once Lupron was discontinued, treated animals began to experience menarche and by the time of data collection for this study, all subjects had reached menarche.

**Figure 1.**
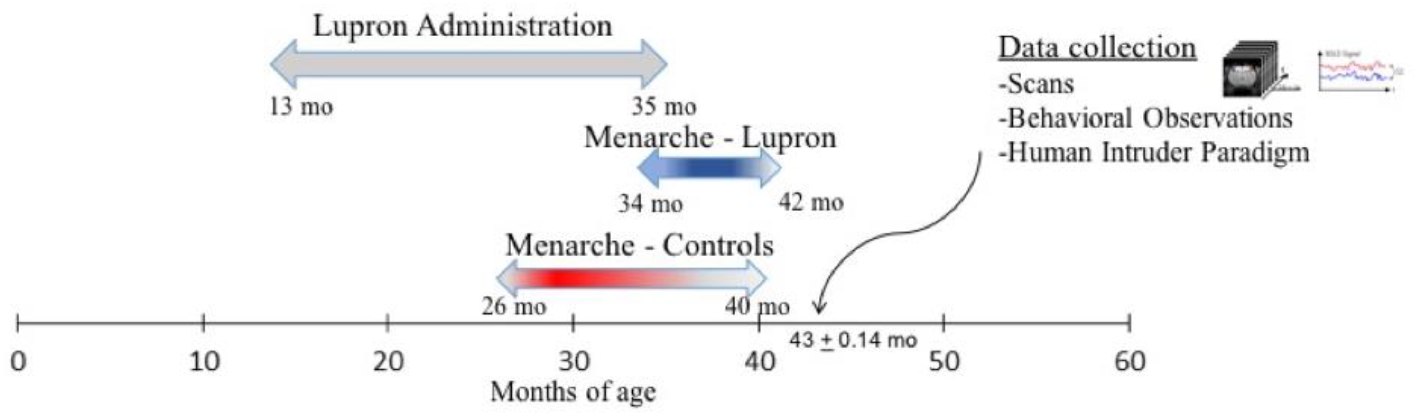
Experimental Timeline. Lupron was administered from prepuberty (16.36±0.23 months of age) through the age typical for menarche for this species (32.56±0.23 months of age). In Control animals, menarche occurred at 30.88±0.64 months of age. Once Lupron was discontinued, Lupron-treated animals began experiencing menarche around 38.07±0.34 months of age. MRI Scans, behavioral observations, and Human Intruder data was collected based on the individual’s age (43.43±0.14 months). All animals had reached menarche by the time of data collection.

### 2.4. Social behavior

Within a month of MRI scanning, seven to eight 30-minute behavioral focal observations were collected for each subject from towers above the social groups using an established rhesus monkey ethogram that includes submissive (withdraw, fear grimace), affiliative (proximity, groom, wrestle/play), anxiety-like (yawn, body shake, self-scratch), and aggressive (threat, display, attack chase) behaviors. Observers recorded the behavior as initiators and recipients in real time with WinObs software. Inter-observer reliability was >92%. One Lupron-treated subject was excluded due to clinical issues and missing data.

### 2.5. Human intruder (HI) task

The HI task measures animals’ emotional reactivity to an unfamiliar human intruder, as we have described previously (Wilson et al., 2013). Sessions were videotaped and established methods were used to score the frequency and duration of behaviors. A Principal Component Analysis (PCA) with varimax rotation was performed using SPSS to reduce the dimensionality of the data. Behaviors with loading scores < 0.4 were excluded from components. Composite scores were calculated for each component and entered as dependent variables in regression models described below. Data were not available for three Lupron-treated subjects.

### 2.6. Functional MRI data acquisition

Resting-state-fMRI and T1 images were acquired at 43.43±0.14 months using a 3T Siemens Tim Trio and an 8-channel phase array coil within the YNPRC Imaging Core. Following induction with telazol (3.81±0.05mg/kg, i.m.), isoflurane anesthesia inhalation was kept to the lowest possible level (1.03±0.01%). Animals were scanned supine; their heads were immobilized in a custom head holder with ear bars and mouthpiece. Four 15-minute fMRI scans were acquired using a T2*-weighted gradient-echo echo-planar imaging (EPI) sequence (400 volumes, TR/TE=2060/25ms, voxel size=1.5mm isotropic). Anatomical scans were acquired using a T1-weighted MPRAGE sequence (128 coronal slices, TR/TE=3000/3.52ms, voxel size=0.5mm isotropic). Subjects were returned to their social group the following day after fully recovering from anesthesia. Imaging data were not collected from one Control.

### 2.7. Functional MRI preprocessing

Imaging data were preprocessed per published protocols (Reding et al., 2019) using an in-house Nipype pipeline incorporating tools from the FMRIB Software Library (FSL; RRID: SCR_002823) and 4dfp tools. The pipeline performs: 1) slice-time correction, 2) one-step resampling of rigid body head motion correction, distortion correction using diffusion field maps, structural-functional co-registration, and non-linear registration of the T1-weighted structural image to the 112RM-SL atlas in F99 space at 1.5×1.5×1.5mm^3^ resolution, 4) signal normalization to a mode of 1000, 5) detrending, 6) regression of rigid body head motion parameters, whole-brain, ventricle, and white matter signal, and all first-order derivatives, and 7) low-pass filtering (< 0.1 Hz). The four runs were concatenated and frames with displacement (FD) greater than 0.2 mm were censored from analysis.

### 2.8. Regions of Interest

For voxelwise analyses, left and right amygdala ROIs were manually drawn using cytoarchitectonic maps in a UNC-Wisconsin adolescent atlas (RRID: SCR_002570), then propagated to the 112 atlas in F99 space with flirt and fnirt tools (FSL; Fig. 5D). For Region of Interest (ROI) analyses, five prefrontal ROIs corresponding to Brodmann Area 46 (BA46 -dorsolateral PFC), BA32 (medial PFC), BA24 (anterior cingulate), BA25 (subgenual cingulate), and BA13 (orbitofrontal cortex) were defined using anatomical parcellations in F99 space (Fig. 5D). Each ROI was then manually edited for neuroanatomical accuracy and to avoid overlap, and masked to avoid regions of signal dropout, using subject-level masks that included voxels exceeding a dropout threshold of the mean intensity of the EPI signal across a whole-brain mask minus two standard deviations.

### 2.9. MRI quality control

One animal with a large lesion, one with a persistent preprocessing fault, nine with substantial EPI artifact, and six animals with biologically implausible patterns in amygdala FC maps (indicative of additional scanning artifacts) were excluded from imaging analyses. Thirteen further subjects with moderate intensity artifact were excluded from voxelwise analyses, while ROI-ROI analyses were performed with and without them. The results section details the specific number of animals included in each analysis.

### 2.10. Resting-state functional connectivity

For voxelwise analyses, the average time course across voxels within each amygdala ROI was extracted and correlated with that of all other brain voxels. Correlation coefficients were Fisher Z-transformed. Using FSL’s FEAT, group-level analyses were performed to identify main and interacting effects of social status (a continuous regressor coding relative social rank) and pubertal delay (Lupron-treated/delayed vs. Control) on whole-brain functional connectivity (FC) within a mask formed by the intersection of 25% gray-matter tissue probability and voxels exceeding a signal dropout threshold (Fig. S1). The native resolution of 1.5mm was maintained to reduce the risk of false positives. Gaussian Random Field-based correction for multiple comparisons was performed (voxel-wise Z>3.1, cluster-level p<0.005, corrected).

For amygdala-PFC ROI-ROI analyses, the average time course across voxels within the right and left amygdala and right and left counterparts of the five prefrontal ROIs described above (BA46, BA32, BA24, BA25, and BA13) was extracted. To limit multiple comparisons, we computed FC between ipsilateral ROI pairs only (i.e., right amygdala-right BA46, left amygdala-left BA13, etc.). The resulting ten measures of amygdala-PFC FC were examined using the same statistical models as the behavioral data, described in the next section.

### 2.11. Statistical analyses

We first verified whether social status (psychosocial stress) and/or pubertal delay affected the timing of pubertal milestones (menarche, first ovulation) using regression models (*lmPerm* in R) that included a continuous regressor coding relative social rank, a binary regressor coding treatment group (Lupron-treated/delayed vs. Control), and an interaction term. Significance was evaluated using a permutation approach and 5000 permutations (perm = “Exact”; fewer than 5000 are performed if the p-value does not approach 0.05).

Next, to examine the effects of subordination stress and experimentally induced pubertal delay on behavior and amygdala-prefrontal FC, we performed a series of regressions. For each dependent variable (i.e., submissive behavior, affiliation (initiating proximity to others), anxiety-like behaviors, aggression received, three principle component scores derived from the Human Intruder task, and ten measures of amygdala-prefrontal FC), relative social rank, treatment condition, and the interaction between social rank and treatment were entered as predictors in the first model (DV = β_0_ + β_1_ Social Rank + β_2_ Treatment + β_3_ Social Rank x Treatment). We did not correct for multiple comparisons, given the relatively small sample size and complex design, but used a robust permutation approach to statistical significance for all tests. Where variables included extreme outliers (less than or greater than 3SD from the mean), analyses were performed with and without outliers.

### 2.12. Brain-behavior analyses

To examine the behavioral significance of FC associations for social status and treatment, we regressed FC between regions (for voxelwise analyses) or nodes (for ROI-ROI analyses) found to exhibit significant effects of social rank or Lupron treatment on the behavioral measures that also showed a significant effect of these factors using the *lm* regression function in R. For each dependent behavioral variable (e.g., affiliative behaviors), we performed a hierarchical regression in which relative social rank, treatment condition, and the interaction between social rank and treatment were entered as predictors in the first step, and FC was entered in the second step. A brain-behavior relationship was considered significant if the addition of FC significantly improved the explanatory power of the model (i.e., the change in R2 was significant, per the *lmSupport* function in R).

Data and scripts for statistical analyses are available from: https://osf.io/8zgrs/

## 3. RESULTS

### 3.1. Pubertal timing

As expected, Lupron-treated animals experienced menarche (*p*<.001) and first ovulation (*p*<.001) significantly later than Controls, confirming the efficacy of the Lupron treatment in delaying these pubertal milestones (Table 1, Fig. 2). There was also an effect of social status, such that more subordinate rank (greater psychosocial stress) was associated with later menarche (*p*=.031) and first ovulation (*p*<.005; Fig. 2) than more dominant rank (lower stress). Significant interactions between social status and Lupron treatment (both p<.01) show that subordinate social status was linearly associated with later menarche and first ovulation in the Control group, but not in the Lupron-treated group (Fig. 2). For Lupron-treated females, menarche and first ovulation occurred within a restricted age range following cessation of Lupron treatment, and showed no relationship with social status. Sixty-nine animals were included in the menarche analyses (Lupron-treated *n*=33; Controls *n*=36). Age of first ovulation was available for 51 animals (Lupron-treated *n*=26; Controls *n*=25).

**Figure 2.**
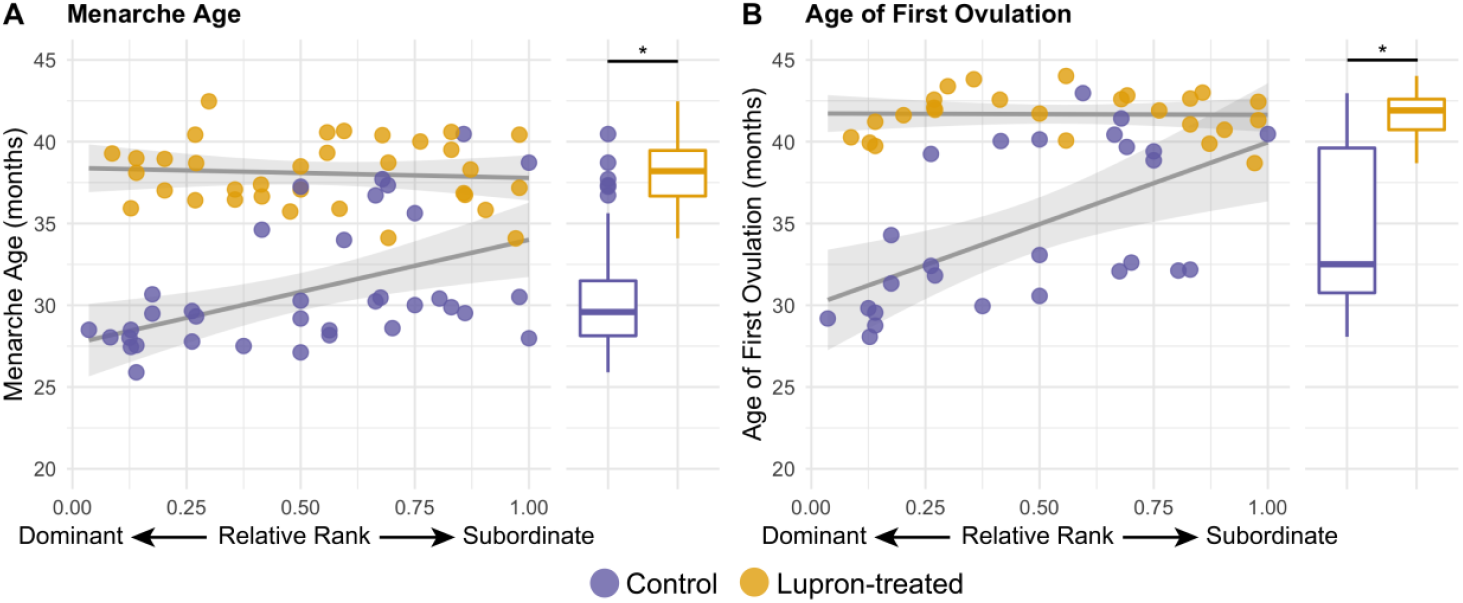
Effect of Lupron treatment and social rank on pubertal timing. **A.** Menarche age was significantly delayed in Lupron-treated subjects relative to Controls. Relative social rank (values close to 1 indicate subordinate status (chronically stressed), while values close to 0 indicate dominant status (unstressed)) was positively associated with menarche age for the Control group, but not for the Lupron-treated group (Table 1). **B.** Lupron-treated subjects experienced first ovulation at a significantly later age than Controls. A positive linear relationship between relative rank and age at first ovulation is evident only amongst Controls.

**Table 1.** Regression analysis of pubertal timing. Social rank (indexing chronic psychosocial stress) and Lupron treatment were significantly associated with menarche age and first ovulation age, with more subordinate ranking females and animals in the Lupron-treated group experiencing later menarche and first ovulation, relative to dominant females and untreated Controls. A relative social rank x Lupron interaction reflects a strong positive relationship between social rank and pubertal timing amongst Control animals, but not those in the Lupron-treated group (see Fig. 2).

|  |  | 95% Confidence Interval |  |  |  |
| --- | --- | --- | --- | --- | --- |
|  | Estimate | SE | LL | UL | *p* |
| First Menstruation |  |  |  |  |  |
| Social Rank | 2.99 | 1.19 | 0.61 | 5.38 | 0.025 |
| Treatment | 7.12 | 0.67 | 5.77 | 8.46 | <.001 |
| Interaction | -7.02 | 2.39 | -11.79 | -2.25 | 0.007 |
| First Ovulation |  |  |  |  |  |
| Social Rank | 5.06 | 1.49 | 2.06 | 8.06 | 0.004 |
| Treatment | 6.69 | 0.86 | 4.96 | 8.41 | <.001 |
| Interaction | -10.08 | 2.98 | -16.07 | -4.09 | 0.005 |

### 3.2. Social behavior

As expected, more subordinate rank was associated with significantly more frequent submissive behaviors toward non-kin (*p*<.001; Table 2; Fig. 3A). This result illustrates the linear dosing effect of social status, with more subordinate females within the hierarchy exhibiting more frequent submissive behaviors. Importantly, however, there was also a significant effect of treatment, with Lupron-treated animals engaging in submissive behaviors more frequently than Controls (*p*<.001). There was no interaction between social status and treatment on the frequency of submissive behaviors (p=0.56).

**Figure 3.**
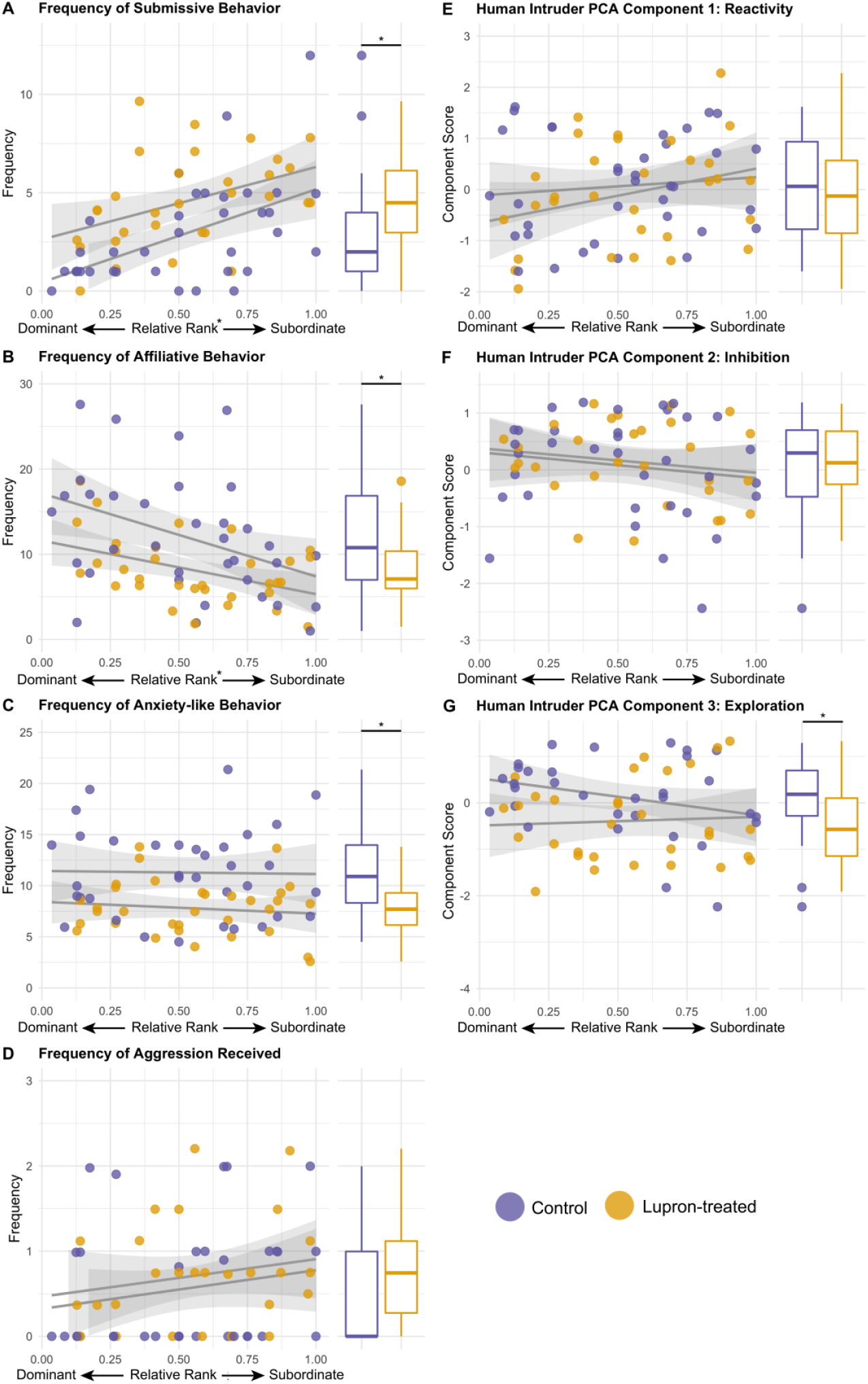
Associations between subordination stress (relative social rank), Lupron treatment, and socioemotional behaviors. **A.** Rate of submissive behaviors toward non-kin was predicted by social rank and Lupron treatment, with more subordinate animals more frequently exhibiting submissive behaviors, relative to dominant animals, and Lupron-treated animals more frequently exhibiting submissive behaviors than Controls. **B.** Rates of proximity initiated with non-kin was negatively predicted by both social rank and Lupron treatment. **C.** Lupron-treated subjects displayed lower anxiety-like behavior in their social groups. **D.** There were no significant effects of social rank or Lupron treatment on aggression received. There were no significant effects of social rank or Lupron treatment on Components 1 (**E**) and 2 (**F**) of the Human Intruder paradigm behaviors. For Component 3 (*Exploration*), Lupron-treated subjects had lower scores than Controls **(G)**. Extreme outliers (</> 3SD from the mean) were excluded from all plots. *Significant at p<.05 – see Table 2.

**Table 2.** Hierarchical regressions: socioemotional behaviors. Regression models examined effects of relative social rank, Lupron treatment, and the interaction between relative social rank and Lupron treatment on socioemotional behavior and behavior during the Human Intruder (HI) task.

|  |  | 95% Confidence Interval |  |  |  |
| --- | --- | --- | --- | --- | --- |
|  | Estimate | SE | LL | UL | p |
| **Submissive Behavior** |  |  |  |  |  |
| Social Rank | 5.32 | 1.11 | 3.10 | 7.54 | <.001 |
| Treatment | 2.10 | 0.62 | 0.85 | 3.34 | <.001 |
| Interaction | 1.18 | 2.23 | -3.28 | 5.63 | 0.562 |
| **Affiliative Behavior** |  |  |  |  |  |
| Social Rank | -10.04 | 2.73 | -15.50 | -4.58 | <.001 |
| Treatment | -4.44 | 1.53 | -7.49 | -1.38 | 0.002 |
| Interaction | 7.16 | 5.49 | -3.80 | 18.12 | 0.114 |
| **Anxiety-like Behavior** |  |  |  |  |  |
| Social Rank | -2.06 | 2.28 | -6.61 | 2.49 | 0.244 |
| Treatment | -4.29 | 1.27 | -6.83 | -1.75 | <.001 |
| Interaction | 1.75 | 4.57 | -7.38 | 10.88 | 0.902 |
| **Aggression Received** |  |  |  |  |  |
| Social Rank | 0.53 | 0.52 | -0.51 | 1.57 | 0.521 |
| Treatment | 0.53 | 0.29 | -0.06 | 1.11 | 0.745 |
| Interaction | 0.29 | 1.05 | -1.80 | 2.38 | 0.706 |
| **HI Component 1: Reactivity** |  |  |  |  |  |
| Social Rank | 0.68 | 0.43 | -0.18 | 1.54 | 0.062 |
| Treatment | -0.17 | 0.24 | -0.65 | 0.32 | 0.725 |
| Interaction | 0.71 | 0.87 | -1.02 | 2.44 | 0.290 |
| **HI Component 2: Inhibition** |  |  |  |  |  |
| Social Rank | 0.02 | 0.44 | -0.86 | 0.91 | 1.000 |
| Treatment | 0.04 | 0.25 | -0.46 | 0.54 | 0.583 |
| Interaction | 0.18 | 0.89 | -1.59 | 1.96 | 0.882 |
| **HI Component 3: Exploration** |  |  |  |  |  |
| Social Rank | -0.24 | 0.41 | -1.06 | 0.57 | 0.430 |
| Treatment | -0.72 | 0.23 | -1.18 | -0.25 | 0.004 |
| Interaction | 0.81 | 0.82 | -0.83 | 2.45 | 0.143 |

Affiliative behaviors also exhibited an effect of social rank, such that more subordinate animals initiated proximity with non-kin less frequently than dominant monkeys (*p*<.001; Table 2; Fig. 3B). There was a significant effect of treatment on affiliative behaviors, with Lupron-treated animals engaging in affiliative behaviors less frequently than Controls (*p*=.002). There was no interaction between social rank and treatment for the frequency of affiliative behaviors (p = 0.11).

There was no significant relationship between social rank and anxiety-like behavior (p=.24; Table 2; Fig. 3C), but there was a significant effect of treatment, with Lupron-treated subjects displaying less frequent anxiety-like behaviors than Controls, (*p*<.001). No interaction was observed.

No main or interaction effects of social rank or Lupron treatment were observed for the frequency of aggression received (Fig. 3D; Relative Rank: *p*=.51; Lupron treatment: *p*=.75; Interaction: *p*=.7). Sixty-nine animals were included in these analyses (Controls *n*=36; Lupron-treated *n*=33). Excluding extreme outliers (less than or greater than 3SD from the mean) did not change these results (Table S1).

### 3.3. HI task: emotional reactivity

Three components were identified in the PCA of HI behaviors and were labeled *Reactivity* (31.43 % of the variance; positive loadings from avert gaze, anxiety-like behaviors, threat, appease, and locomote), *Behavioral Inhibition* (22.07%; positive loadings from freezing, negative loading from locomotion), and *Exploration* (17.51%; positive loadings from exploration and negative from freezing).

No main or interacting effects were obtained for the first two components, *Reactivity* and *Behavioral Inhibition*, although an association between social status and Reactivity was just outside the significance threshold (p=.062; Table 2; Fig. 3E). Lupron treatment was associated with lower *Exploration* scores, but there was no effect of social status (*p*=.004; Table 2; Fig. 3G). Sixty-nine animals were included in these analyses (Controls *n*=36; Lupron-treated *n*=33). Excluding extreme outliers (less than or greater than 3SD from the mean) did not change the results (Table S1).

### 3.4. Voxelwise resting-state Functional Connectivity (FC)

Group-level voxelwise FC of amygdala across all subjects (*n*=39; Controls *n*=19; Lupron-treated *n*=20) encompassed a bilateral network including hippocampus, temporal pole, superior temporal sulcus and gyrus, insula, and dorsal anterior cingulate (Fig. 4A). At a stringent statistical threshold (Z>3.1; p<.005), there was a significant positive association between relative social rank and FC between left amygdala and right temporal pole, such that more subordinate status was associated with stronger FC (cluster volume=148.5mm^3^; Fig. 4B). At a less stringent statistical threshold (Z>2.3; p<.005), the extent of the positive association between social rank and FC between left amygdala and right temporal pole was increased (cluster volume=556.88mm^3^), and there was a further positive association between social rank and FC between right amygdala and left ventrolateral PFC (BA 47/12; cluster volume=33.75mm^3^; Fig. S2). There were no significant effects of treatment on voxelwise FC.

**Figure 4.**
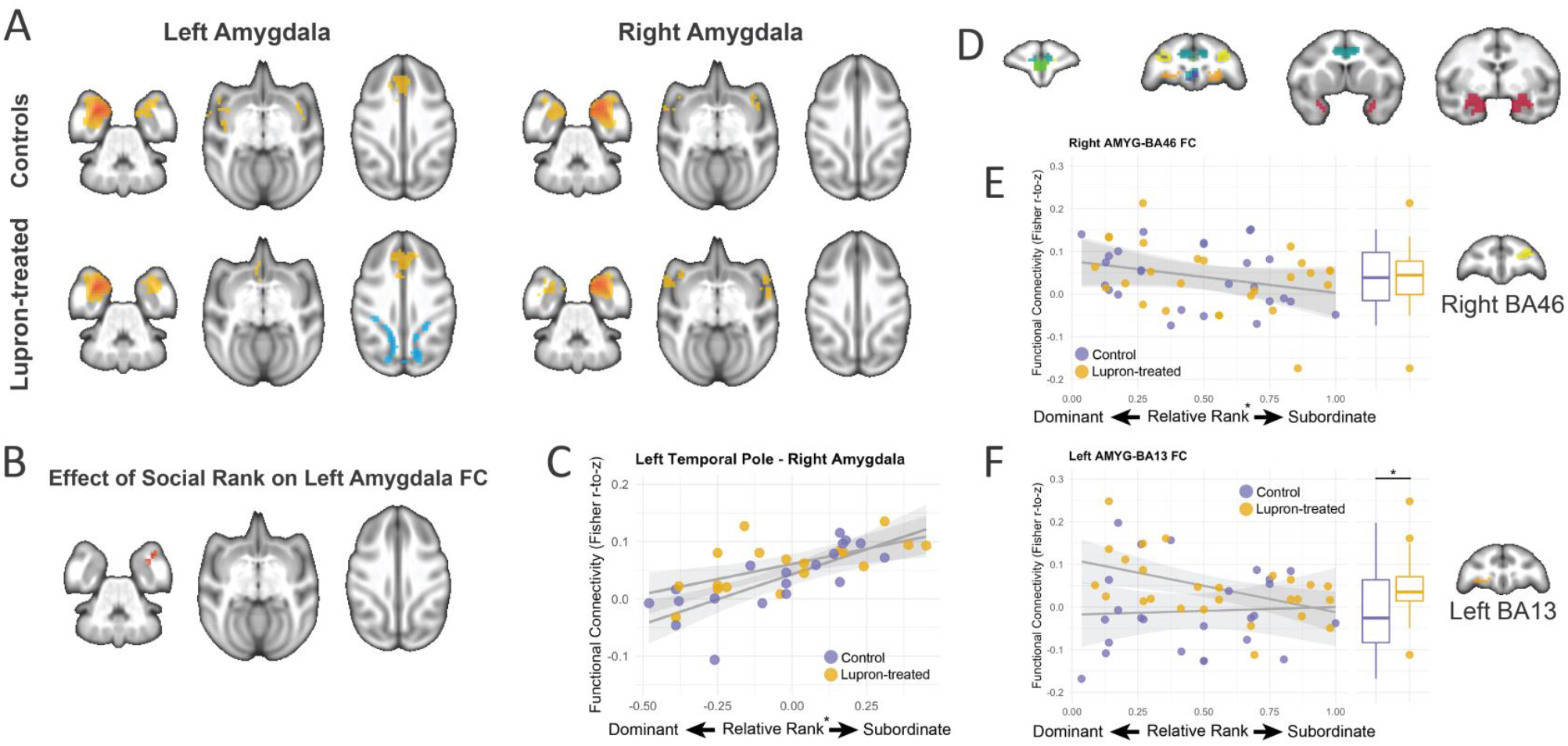
Amygdala Functional Connectivity is altered by social rank and pubertal delay. **A.** Voxelwise functional connectivity of left and right amygdala (ROIs shown in D) in the Control and Lupron-treated groups. **B.** Positive association between social status (relative rank, indexing exposure to chronic stress) and FC between left amygdala and right temporal pole; the scatterplot in **C** shows that more subordinate monkeys (higher relative rank) showed stronger FC between the left amygdala ROI and right temporal pole. **D**. Anatomically defined Regions of Interest (ROIs) located in amygdala (red), BA46 (yellow), BA32 (green), BA24 (teal), BA25 (purple), and BA13 (orange). **E.** FC between right amygdala and right dorsolateral PFC (BA46) was negatively associated with subordination stress, with more subordinate monkeys showing weaker FC than dominant ones (Table 4; Model II). **F.** Pubertal delay was associated with FC between left amygdala and left orbitofrontal cortex (BA13) (Table 4), such that Lupron-treated animals exhibited stronger FC than Controls.

### 3.5. ROI-ROI analyses

Targeted amygdala-PFC ROI-ROI analyses (*n*=52; Controls *n*=26; Lupron-treated *n*=26) revealed two significant associations. First, FC between right amygdala and right dlPFC (BA46) was negatively associated with social rank, with more subordinate monkeys showing weaker FC than dominant ones (*b*=-0.074, SE = 0.036, 95 CI = [-0.146, - 0.002], *p* = 0.029; Table S2; Fig. 4E). Second, Lupron treatment was associated with FC between left amygdala and left orbitofrontal cortex (BA13) (*b* = 0.067, SE = 0.025, 95 CI = [0.018, 0.117], *p* = 0.014; Table S2; Fig. 4F), such that Lupron-treated animals exhibited stronger FC than Controls in this circuit.

No other effects of social rank or Lupron treatment were observed (all *p*s>.062; Table S2; Fig S3). Excluding 13 subjects with mild artifact in their resting-state fMRI scans did not change these two findings, although a significant rank x treatment interaction was observed for FC between right amygdala and BA32 (p=.03; Table S3).

### 3.6. Brain-Behavior Relationships

To minimize multiple comparisons, we restricted analyses of brain-behavior relationships to functional connections that exhibited significant effects of social rank or Lupron treatment and behavioral measures that also showed significant effects of social rank or treatment. Accordingly, 12 brain-behavior relationships were examined: between three functional connections (left amygdala-right temporal pole (mean FC within the significant cluster identified in voxelwise analyses, i.e., Figure 4B), right amygdala and right dlPFC (BA46), and left amygdala and left orbitofrontal cortex (BA13)) and four behaviors (frequency of submissive, affiliative, and anxiety-like behaviors, and scores on HI Component 3: Exploration).

When outliers (</> 3SD from the mean) were excluded, none of the brain-behavior relationships examined were significant, although there were two brain-behavior relationships that were just above the threshold for significance - for voxelwise FC between left amygdala and temporal and affiliative behaviors (*n*=37; *b*=0.0021, SE = 0.001, 95 CI = [-0.000025, 0.0041], p = 0.053) and Human Intruder Exploration component scores (*n*=36; *b*=-0.013, SE = 0.0064, 95 CI = [-0.0262, 0.00007], p = 0.052).

## 4. DISCUSSION

Our goal was to examine the effect of the timing of the pubertal rise in gonadal hormones (typical or experimentally delayed) on neurobehavioral correlates of chronic psychosocial stress (subordinate social status) in adolescent female rhesus macaques. Specifically, we sought to test opposing hypotheses – (1) that subordination stress would have robust negative effects, regardless of chronological age at puberty onset (i.e., there would be no interaction between social status and pubertal delay), and (2) experimentally delaying puberty would protect against the effects of subordination stress on neurobehavioral outcomes (i.e., social status and pubertal delay would interact, such that the impact of subordination stress would be ameliorated in *treated* subordinate monkeys, whose puberty was experimentally delayed). Our findings support the first alternative - although Lupron treatment abolished the known effect of social subordination stress on pubertal onset (i.e., later age of menarche and first ovulation), subordinate status (greater stress) was associated with altered socioemotional behavior and amygdala FC, regardless of treatment (i.e., regardless of whether puberty onset was typical or delayed). No interactions between social rank and experimentally induced pubertal delay through treatment with Lupron were detected for any neurobehavioral outcomes. Further, we observed several main, non-interacting, effects of experimentally induced pubertal delay via Lupron treatment on behavior and FC that offer further support for the idea that exposure to puberty-related increases in female gonadal hormones increases vulnerability to socioemotional dysregulation, even when such exposures occur at a later chronological age. These findings highlight a need for further investigations of the interacting impact of pubertal hormones and stress on neurodevelopment, as discussed in more detail below.

### 4.1. Experimental and stress-related delay of puberty in female group-housed macaques

Lupron treatment was effective in delaying the onset of puberty, evidenced by the delayed age of menarche and first ovulation in Lupron-treated animals. An interaction between social rank and Lupron-induced pubertal delay was also detected, such that the well-established linear association between social status and menarche age and first ovulation observed in untreated animals (Wilson et al., 2013) was abolished in Lupron-treated animals. As previously suggested by Wilson et al. (Wilson et al., 2004), it is likely the HPG axis is primed for puberty onset at a specific age for each animal, potentially under the control of a biological clock or critical body weight. Under Lupron treatment, puberty onset was suppressed, but unfolded relatively quickly after Lupron treatment was discontinued, masking any relationship between social rank and pubertal timing.

### 4.2. Social status and experimental pubertal delay separately affect behavior

We obtained a number of expected associations between social status and behavior in the social group (Fig. 3) as well as in response to a standardized behavioral test of a threatening stimulus (Human Intruder task; Fig. 4). The positive association between social rank and frequency of submissive behaviors (Fig. 3A) confirms the hallmark feature of the social subordination model (e.g., Michopoulos et al., 2012). Subordinate-ranking monkeys also initiated proximity with non-kin less frequently than dominant-ranking monkeys (Fig. 3B), consistent with observations that subordinate monkeys engage less in affiliative social behavior (Michopoulos et al., 2012; Snyder-Mackler et al., 2016). Affiliative behavior (e.g., seeking proximity, grooming) serves an important social bonding function amongst unrelated group-living female macaques, and the relatively reduced opportunities for prosocial interactions experienced by subordinate-ranking animals may be a critical factor that accounts for the adverse effects of chronic stress on neurobehavioral and health related outcomes (Howell et al., 2014; Reding et al., 2019; Sapolsky, 2005; Snyder-Mackler et al., 2016).

Rates of anxiety-like behaviors, aggression received, and behavioral responses during the Human Intruder Task did not differ in association with social status, however. This might seem inconsistent with the well-established observation that subordinate animals receive more aggression from group mates (Wilson, 2016) and are stressed and therefore expected to be more anxious and behaviorally inhibited. While overwhelming data suggest anxiety is one of many stress-related phenotypes in human adolescents, the expression of anxiety-like behavior may not result from subordinate status in monkeys, particularly for animals living in complex social groups. In fact, no clear relationship between rank in the social hierarchy and anxiety-like behaviors has been established (Michopoulos et al., 2012; Wilson et al., 2013). As noted in the results section, an association between social status and the first principal component of the Human Intruder task fell just above the threshold for significance, suggesting a weak relationship between subordinate status and more reactive behavior (gaze-aversion, anxiety-like behaviors, appeasing behavior, and movement) on the task. Finally, the absence of an association between aggression received and social status may reflect the fact that subordinate animals successfully reduced the probability of receiving aggression by moving away from dominant animals (i.e., they showed reduced affiliative social behavior with dominants), a feature of submissive behavior.

No interactions between social rank and Lupron treatment were detected for any behavioral outcome - pubertal delay did not appear to ameliorate the effects of subordination stress on neurobehavioral outcomes. Yet, several main effects of treatment (pubertal delay) were found that were similar to the effects of subordinate status, challenging the hypothesis that delayed exposure to puberty-linked increases in female gonadal hormones would be protective. Instead, late-onset puberty led to stress-related phenotypes, regardless of social rank. Specifically, like untreated subordinate animals who experienced typical puberty, Lupron-treated animals were more submissive than untreated animals, and importantly, they were also less affiliative and exhibited lower scores on the Exploration component in the HI paradigm (i.e., they explored less and froze more), regardless of their social status. Thus, the rise in gonadal hormones experienced by Lupron-treated animals, who experienced menarche at later ages, and more recently (~5 months) than Control animals (~12 months), relative to the timing of data collection, may have conferred increased vulnerability to socioemotional dysregulation, regardless of social status. Alternatively, the similarities in behavioral outcomes between untreated subordinates and treated animals may be due to the common factor of delayed puberty – whether this is stress-related (untreated animals) or pharmacologically-induced (treated animals). It is thought that exposure to elevated gonadal hormones is a necessary trigger for neuroplasticity supporting developmentally appropriate learning during an adolescent critical period (e.g., Dahl et al., 2018). In Lupron-treated animals, blocking this individual biological trigger, even temporarily, may have enduringly disrupted the maturation of adult behaviors and capacities, since the critical window may have ended or been substantially narrower due to the delay in puberty. In this way, the behavioral effects of Lupron treatment we obtained are similar to previous reports of an association between ovariectomy and reduced affiliative behavior (e.g., Coleman et al., 2011).

Lupron treatment was also associated with lower rates of anxiety-like behavior, while there was no association between subordinate status and anxiety. As stated above, there is no clear relationship between rank in the social hierarchy and anxiety-like behaviors (Michopoulos et al., 2012; Wilson et al., 2013). Michopoulos et al. (2012) found that E2 replacement in ovariectomized macaques significantly reduced anxiety-like behavior; the more recent onset of puberty in the Lupron-treated group (and the associated rise in E2) may therefore explain lower anxiety amongst Lupron-treated animals, relative to Controls. On the other hand, the effects of E2 on anxiety may be context dependent (e.g., Morgan et al., 2004) and the notion that E2 is anxiolytic vs. anxiogenic is likely an oversimplification that ignores the social context.

### 4.3. Social subordination and experimental pubertal delay separately affect functional connectivity

The overall pattern of amygdala FC (Fig. 5A) was consistent with previous FC studies (e.g., Grayson et al., 2016; Reding et al., 2019) and the known anatomical connectivity of macaque amygdala (Höistad and Barbas, 2008), which is reciprocally interconnected with medial temporal pole, hippocampus, parahippocampal cortex, anterior insula, posterior orbitofrontal and medial PFC. In voxelwise, whole-brain analyses with stringent correction for multiple comparisons, we observed only one significant effect – subordinate status was associated with increased FC between left amygdala and right temporal pole (Fig. 5B). The temporal pole is a polymodal association area (Sallet et al., 2011) that has been centrally implicated in social and emotional processing in both humans and monkeys. For example, in an innovative study, Sliwa and Freiwald (Bickart et al., 2012) collected fMRI data while adult male macaque monkeys viewed videos of social interactions between conspecifics. They identified a social interaction processing network that included medial and dorsomedial prefrontal cortices, ventrolateral PFC and temporal pole that they argue supports the analysis of social interactions, potentially an evolutionary precursor of human theory of mind. Adult male macaques with larger social networks have greater grey matter volumes in a network that included amygdala and temporal pole (Bickart et al., 2012), and humans with larger social networks have stronger FC within amygdala circuits that also include temporal pole (Bickart et al., 2012), consistent with a role for these circuits in processing social cues and integrating social information with motivation and emotion to promote social affiliation (Bickart et al., 2012). The increased amygdala-temporal pole FC we observed in subordinate animals may therefore reflect a neurocognitive adaptation to optimize processing of relevant social stimuli. We previously observed similar, potentially adaptive effects of social rank on white matter integrity in prepubertal female macaques, using diffusion tensor imaging (Howell et al., 2014). These observations are consistent with the idea that the subordinate phenotype does not necessarily represent a pathological condition; instead, the experience of being subordinate likely leads to behavioral or physiological adaptations that enable subordinate animals to successfully navigate their social environments and minimize the risk of aggression from more dominant animals (Wilson, 2016).

We were somewhat surprised not to see other significant effects of subordination stress, Lupron treatment, or their interaction at the whole-brain level, but this may be due, in part, to our sample size and application of appropriately stringent corrections for multiple comparisons. Targeted amygdala-PFC ROI-ROI FC analyses revealed only two further main effects. First, we found that FC between right amygdala and right dorsolateral PFC (BA46) was associated with relative rank, such that FC was weaker in subordinate animals, relative to dominants. Second, we found a main effect of Lupron treatment on FC between left amygdala and left orbitofrontal cortex (OFC; BA13), with Lupron-treated animals exhibiting stronger FC than Controls. In the absence of significant brain-behavior relationships, inferences about the behavioral significance of our FC findings are speculative. Identifying the nature of the individual differences in brain organization that confer resilience and vulnerability is a key piece of the psychopathology puzzle, however. For example, human adolescents at risk for depression in whom depression did not develop (“resilient” individuals) exhibited stronger connectivity between amygdala and PFC (dorsolateral and orbitofrontal) than at-risk adolescents in whom depression did develop (“converted” individuals) (Fischer et al., 2018). Our data suggest that while stress and the timing of puberty-linked changes in gonadal hormones may have some overlapping behavioral effects, their effects may be distinguishable at the level of the brain. Fully uncovering the nature of these effects and their long-term behavioral significance may ideally be investigated in a larger and statistically well-powered longitudinal study, potentially more feasible in rodents than in non-human primates.

### 4.4. Limitations

In addition to the small sample size, which was likely underpowered to detect some of the interacting effects hypothesized, a primary limitation of this study is that MRI data collection was conducted at only one time-point that was fixed based on age. Larger-scale fully-powered longitudinal studies featuring multiple data points before, immediately following menarche, and at a first time point post menarche, would precisely control for the duration of E2 exposure independent of chronological age. A related point is that we cannot determine whether neurobehavioral effects of subordinate status are related to the monkeys’ postnatal experience in the social group, or to prenatal factors including maternal gestational stress or inherited genetic/epigenetic differences related to social rank. Future studies taking a lifespan approach beginning prenatally may be better positioned to disentangle these possibilities.

Second, our imaging data revealed fewer effects than expected, particularly in the context of the robust behavioral effects of both social rank and experimental delay of puberty. It is possible that issues like image quality (our data exhibited several imaging artifacts necessitating the exclusion of a number of subjects) and depth of anesthesia may have negatively impacted our ability to detect effects. Advances in MRI data acquisition (for both humans and animals) and the use of sedatives such as medetomidine over anesthesia may produce more robust data in future studies.

## 5. CONCLUSIONS

The striking sex disparity in vulnerability to stress-linked disorders such as anxiety and depression begins at the onset of puberty and is evident throughout the reproductive life cycle (Thapar et al., 2012), suggesting adverse experiences and psychosocial factors interact with female biology to increase vulnerability to behavioral disorders. We do not fully know why this happens, and animal studies have historically tended to focus on adult male animals (Shansky and Woolley, 2016), thus neglecting the role of both sex and development in vulnerability to psychopathology. Here, using an ethologically valid and translational female macaque model of chronic psychosocial stress, we sought to assess whether female pubertal hormones exacerbate the effects of chronic stress on behavior and brain during adolescence by experimentally manipulating the timing of puberty. We observed several effects of subordinate social status on a number of behavioral outcomes. Lupron-induced pubertal delay produced a similar behavioral phenotype to subordinate social status. At the level of the brain, however, the effects of social status and experimental pubertal delay were distinct. Taken together, our findings suggest that while some behavioral phenotypes show similarities in their sensitivity to the effects of stress and to the timing of pubertal hormones, these effects may be distinguishable at the level of the brain. Future longitudinal studies are needed to determine precisely how the timing of E2 exposure during the transition through adolescence alters brain structure and function linked to specific cognitive, emotional, and social behavioral outcomes and how these relationships are modulated by exposure to adverse experiences.

## ACKNOWLEDGEMENTS

The authors would like to thank Jennifer Whitely, Shannon Moss, Angela Tripp, Marta Checchi, Erin O’Sheil, Christine Marsteller, and Natalie Brutto for their exceptional technical contributions. They also thank the staff at the YNPRC for their dedicated animal care and its Imaging Core for excellent services. The YNPRC is fully accredited by AAALAC International.

## 9. SUPPLEMENTAL TABLES

**Table S1.** Hierarchical regressions: socioemotional behaviors, with extreme outliers (greater than or less than 3SD from the mean) removed. Regression models examined effects of relative social rank, Lupron treatment, and the interaction between relative social rank and Lupron treatment on socioemotional behavior and behavior during the Human Intruder (HI) task. Note that for the HI task, only Component 2 exhibited extreme outliers.

|  |  | 95% Confidence Interval |  |  |  |
| --- | --- | --- | --- | --- | --- |
|  | Estimate | SE | LL | UL | p |
| **Submissive Behavior** |  |  |  |  |  |
| Social Rank | 4.26 | 0.97 | 2.33 | 6.20 | <.001 |
| Treatment | 1.64 | 0.54 | 0.56 | 2.71 | 0.003 |
| Interaction | -1.06 | 1.96 | -4.97 | 2.85 | 0.745 |
| **Affiliative Behavior** |  |  |  |  |  |
| Social Rank | -8.09 | 2.34 | -12.77 | -3.40 | <.001 |
| Treatment | -3.69 | 1.30 | -6.29 | -1.09 | 0.011 |
| Interaction | 3.47 | 4.70 | -5.91 | 12.85 | 0.433 |
| **Anxiety-like Behavior** |  |  |  |  |  |
| Social Rank | -0.69 | 1.61 | -3.90 | 2.52 | 0.824 |
| Treatment | -3.47 | 0.90 | -5.27 | -1.68 | <.001 |
| Interaction | -0.88 | 3.22 | -7.32 | 5.56 | 0.667 |
| **Aggression Received** |  |  |  |  |  |
| Social Rank | 0.31 | 0.31 | -0.31 | 0.93 | 0.824 |
| Treatment | 0.19 | 0.17 | -0.16 | 0.54 | 0.161 |
| Interaction | -0.18 | 0.63 | -1.43 | 1.07 | 0.824 |
| **HI Component 2: Inhibition** |  |  |  |  |  |
| Social Rank | 0.02 | 0.44 | -0.86 | 0.91 | 1.000 |
| Treatment | 0.04 | 0.25 | -0.46 | 0.54 | 0.583 |
| Interaction | 0.18 | 0.89 | -1.59 | 1.96 | 0.882 |

**Table S2.** Hierarchical regressions: ROI-ROI relationships. Regression models examined effects of relative social rank, Lupron treatment, and the interaction between relative social rank and Lupron treatment on functional connectivity between the left and right amygdala and five prefrontal ROIs corresponding to BA46, BA32, BA24, BA25, and BA13 (Fig. 5D).

|  |  | 95% Confidence Interval |  |  |  |
| --- | --- | --- | --- | --- | --- |
|  | Estimate | SE | LL | UL | p |
| Left Amyg-BA46 |  |  |  |  |  |
| Social Rank | 0.05 | 0.04 | -0.03 | 0.14 | 0.548 |
| Treatment | -0.01 | 0.02 | -0.06 | 0.04 | 0.516 |
| Interaction | -0.05 | 0.08 | -0.22 | 0.12 | 0.619 |
| Right Amyg-BA46 |  |  |  |  |  |
| Social Rank | -0.07 | 0.04 | -0.15 | 0.00 | 0.029 |
| Treatment | 0.00 | 0.02 | -0.04 | 0.04 | 0.980 |
| Interaction | 0.00 | 0.07 | -0.14 | 0.14 | 1.000 |
| Left Amyg-BA32 |  |  |  |  |  |
| Social Rank | 0.04 | 0.04 | -0.03 | 0.12 | 0.250 |
| Treatment | 0.01 | 0.02 | -0.03 | 0.05 | 0.980 |
| Interaction | -0.02 | 0.07 | -0.17 | 0.13 | 1.000 |
| Right Amyg-BA32 |  |  |  |  |  |
| Social Rank | -0.03 | 0.04 | -0.12 | 0.06 | 0.725 |
| Treatment | 0.02 | 0.02 | -0.03 | 0.07 | 0.643 |
| Interaction | -0.05 | 0.09 | -0.22 | 0.12 | 0.686 |
| Left Amyg-BA24 |  |  |  |  |  |
| Social Rank | -0.03 | 0.04 | -0.10 | 0.04 | 0.362 |
| Treatment | -0.02 | 0.02 | -0.06 | 0.02 | 0.765 |
| Interaction | 0.09 | 0.07 | -0.06 | 0.24 | 0.168 |
| Right Amyg-BA24 |  |  |  |  |  |
| Social Rank | -0.02 | 0.05 | -0.11 | 0.07 | 0.980 |
| Treatment | -0.02 | 0.03 | -0.07 | 0.04 | 0.338 |
| Interaction | 0.04 | 0.09 | -0.14 | 0.22 | 0.521 |
| Left Amyg-BA25 |  |  |  |  |  |
| Social Rank | 0.00 | 0.04 | -0.07 | 0.08 | 0.941 |
| Treatment | 0.04 | 0.02 | 0.00 | 0.08 | 0.062 |
| Interaction | -0.06 | 0.07 | -0.21 | 0.09 | 0.394 |
| **Right Amyg-BA25** |  |  |  |  |  |
| Social Rank | 0.01 | 0.04 | -0.07 | 0.09 | 0.765 |
| Treatment | 0.00 | 0.02 | -0.05 | 0.04 | 1.000 |
| Interaction | 0.05 | 0.08 | -0.11 | 0.22 | 0.420 |
| **Left Amyg-BA13** |  |  |  |  |  |
| Social Rank | -0.06 | 0.04 | -0.15 | 0.02 | 0.435 |
| Treatment | 0.07 | 0.02 | 0.02 | 0.12 | 0.014 |
| Interaction | -0.12 | 0.09 | -0.29 | 0.06 | 0.138 |
| **Right Amyg-BA13** |  |  |  |  |  |
| Social Rank | -0.06 | 0.05 | -0.16 | 0.05 | 0.594 |
| Treatment | 0.03 | 0.03 | -0.03 | 0.09 | 0.225 |
| Interaction | -0.03 | 0.10 | -0.24 | 0.18 | 0.902 |

**Table S3.**
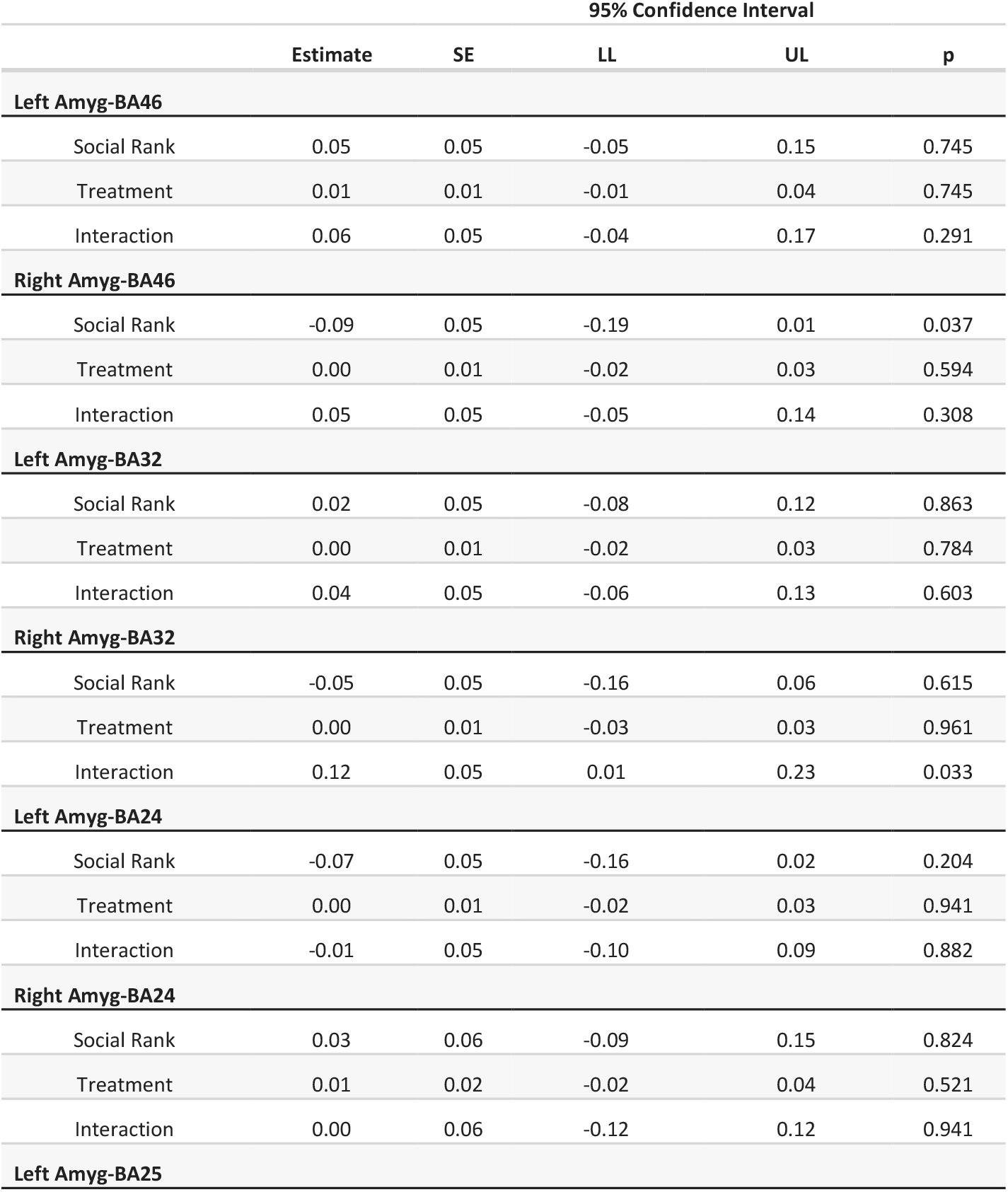
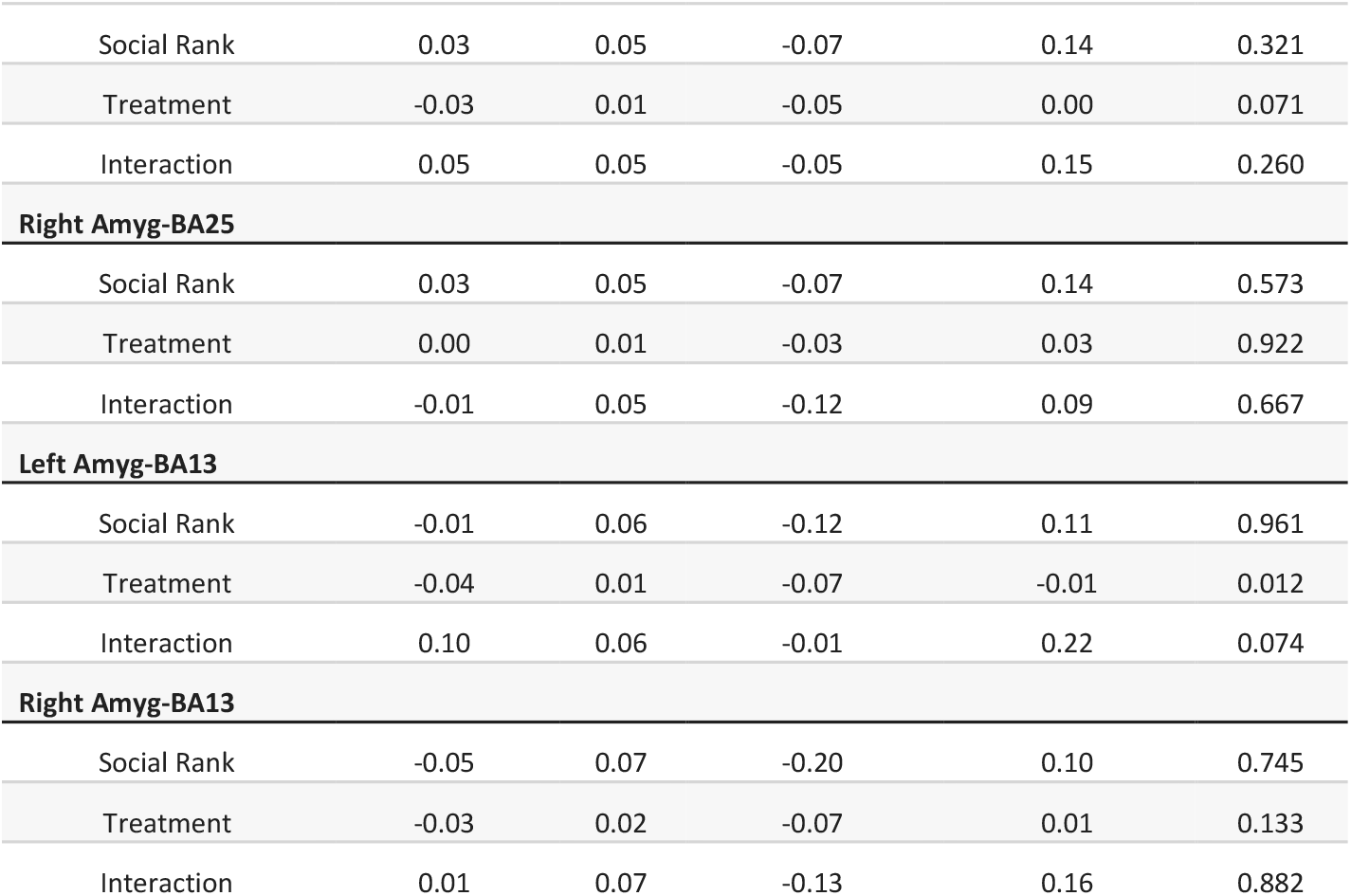
Hierarchical regressions: ROI-ROI relationships, excluding 13 subjects with mild artifact in their resting-state fMRI scans. Regression models examined effects of relative social rank, Lupron treatment, and the interaction between relative social rank and Lupron treatment on functional connectivity between the left and right amygdala and five prefrontal ROIs corresponding to BA46, BA32, BA24, BA25, and BA13 (Fig. 5D).

## 10. SUPPLEMENTAL FIGURES AND CAPTIONS

**Figure S1.**
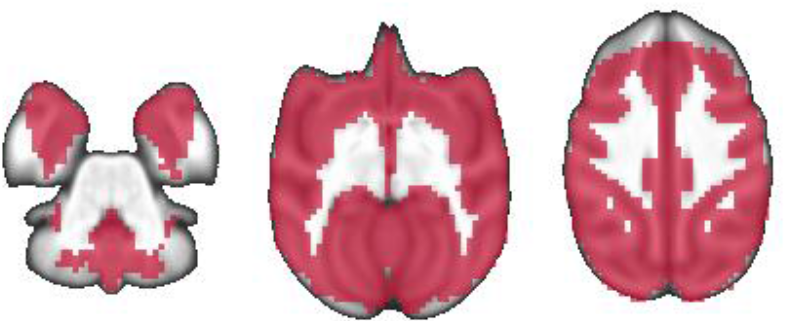
Mask applied in whole-brain functional connectivity analyses. The mask was formed by the intersection of 25% gray-matter tissue probability and voxels exceeding a dropout threshold of the mean intensity of the EPI signal minus two standard deviations across all subjects.

**Figure S2.**
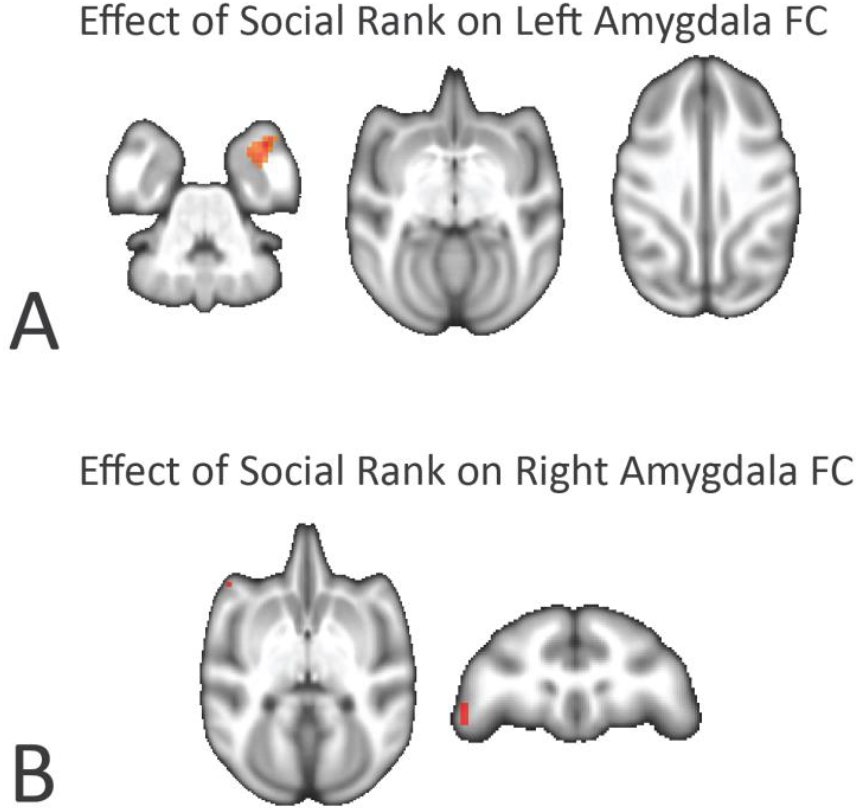
**A.** The extent of the positive association between stress and FC between left amygdala and right temporal pole is increased when a less stringent statistical threshold (Z>2.3; p<0.005) is applied. **B.** At this threshold, a further positive association between stress and FC between right amygdala and left dorsolateral PFC is detected.

**Figure S3.**
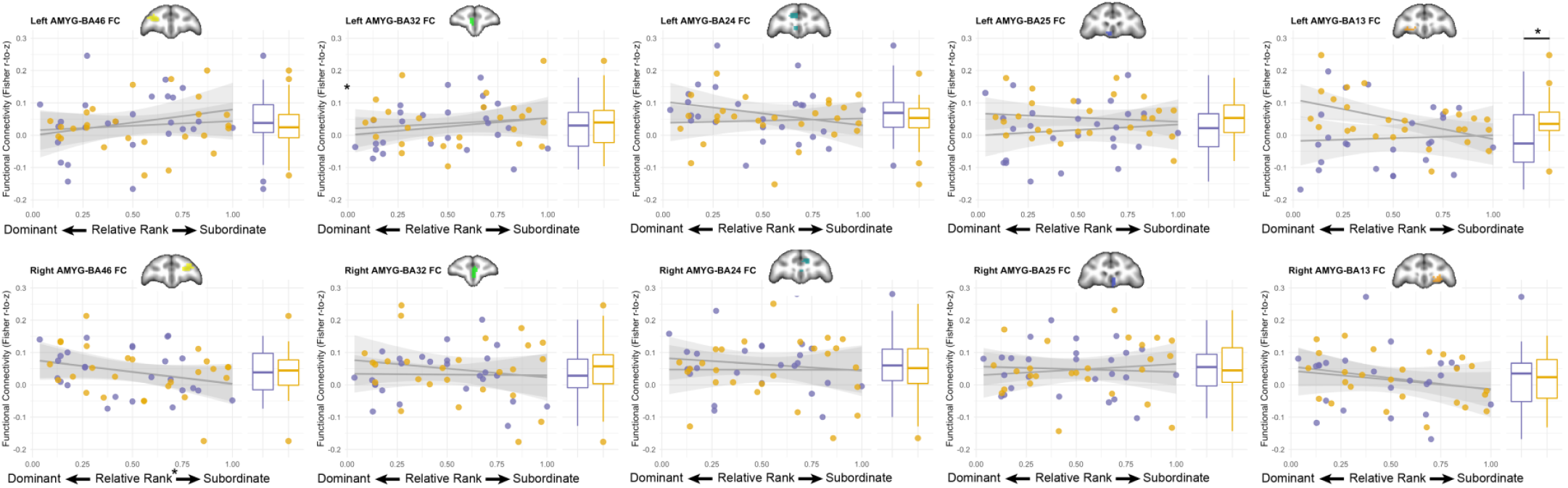
Results of amygdala-PFC ROI analyses. Significant effects of stress (relative rank) and PTL were obtained for FC between right amygdala and right dorsolateral PFC (BA46; **B**) and FC between left amygdala and left orbitofrontal cortex (BA13; **I**), as reported in Table S2 and shown in Fig. 5.

## REFERENCES

Bickart, K.C., Hollenbeck, M.C., Barrett, L.F., Dickerson, B.C., 2012. Intrinsic amygdala-cortical functional connectivity predicts social network size in humans. J Neurosci 32, 14729–14741. doi:10.1523/JNEUROSCI.1599-12.2012

Bourke, C.H., Neigh, G.N., 2011. Behavioral effects of chronic adolescent stress are sustained and sexually dimorphic. Horm Behav 60, 112–120. doi:10.1016/j.yhbeh.2011.03.011

Breach, M.R., Moench, K.M., Wellman, C.L., 2019. Social instability in adolescence differentially alters dendritic morphology in the medial prefrontal cortex and its response to stress in adult male and female rats. Dev Neurobiol 79, 839–856. doi:10.1002/dneu.22723

Cohodes, E.M., Kitt, E.R., Baskin-Sommers, A., Gee, D.G., 2020. Influences of early-life stress on frontolimbic circuitry: Harnessing a dimensional approach to elucidate the effects of heterogeneity in stress exposure. Dev Psychobiol 55, 389. doi:10.1002/dev.21969

Coleman, K., Robertson, N.D., Bethea, C.L., 2011. Long-term ovariectomy alters social and anxious behaviors in semi-free ranging Japanese macaques. Behav Brain Res 225, 317–327. doi:10.1016/j.bbr.2011.07.046

Copeland, W.E., Worthman, C., Shanahan, L., Costello, E.J., Angold, A., 2019. Early Pubertal Timing and Testosterone Associated With Higher Levels of Adolescent Depression in Girls. J Am Acad Child Adolesc Psychiatry 58, 1197–1206. doi:10.1016/j.jaac.2019.02.007

Dahl, R.E., Allen, N.B., Wilbrecht, L., Suleiman, A.B., 2018. Importance of investing in adolescence from a developmental science perspective. Nature 554, 441–450. doi:10.1038/nature25770

Eiland, L., Ramroop, J., Hill, M.N., Manley, J., McEwen, B.S., 2012. Chronic juvenile stress produces corticolimbic dendritic architectural remodeling and modulates emotional behavior in male and female rats. Psychoneuroendocrinology 37, 39–47. doi:10.1016/j.psyneuen.2011.04.015

Eugster, E.A., 2019. Treatment of Central Precocious Puberty. J Endocr Soc 3, 965–972. doi:10.1210/js.2019-00036

Fareri, D.S., Gabard-Durnam, L., Goff, B., Flannery, J., Gee, D.G., Lumian, D.S., Caldera, C., Tottenham, N., 2017. Altered ventral striatal-medial prefrontal cortex resting-state connectivity mediates adolescent social problems after early institutional care. Dev Psychopathol 29, 1865–1876. doi:10.1017/S0954579417001456

Fischer, A.S., Camacho, M.C., Ho, T.C., Whitfield-Gabrieli, S., Gotlib, I.H., 2018. Neural Markers of Resilience in Adolescent Females at Familial Risk for Major Depressive Disorder. JAMA Psychiatry 75, 493–502. doi:10.1001/jamapsychiatry.2017.4516

Fuhrmann, D., Knoll, L.J., Blakemore, S.-J., 2015. Adolescence as a Sensitive Period of Brain Development. Trends Cogn Sci (Regul Ed) 19, 558–566. doi:10.1016/j.tics.2015.07.008

Goddings, A.-L., Beltz, A., Peper, J.S., Crone, E.A., Braams, B.R., 2019. Understanding the Role of Puberty in Structural and Functional Development of the Adolescent Brain. J Res Adolesc 29, 32–53. doi:10.1111/jora.12408

Godfrey, J.R., Pincus, M., Sanchez, M.M., 2016. Effects of Social Subordination on Macaque Neurobehavioral Outcomes: Focus on Neurodevelopment, in: Social Inequalities in Health in Nonhuman Primates, Developments in Primatology: Progress and Prospects. Springer, Cham, Cham, pp. 25–47. doi:10.1007/978-3-319-30872-2_3

Grayson, D.S., Bliss-Moreau, E., Machado, C.J., Bennett, J., Shen, K., Grant, K.A., Fair, D.A., Amaral, D.G., 2016. The Rhesus Monkey Connectome Predicts Disrupted Functional Networks Resulting from Pharmacogenetic Inactivation of the Amygdala. Neuron 91, 453–466. doi:10.1016/j.neuron.2016.06.005

Herman-Giddens, M.E., 2017. Lessons Learned From Australia: Social Disadvantage and Pubertal Timing. Pediatrics 139, e20170837. doi:10.1542/peds.2017-0837

Herman-Giddens, M.E., 2006. Recent data on pubertal milestones in United States children: the secular trend toward earlier development. Int. J. Androl. 29, 241–6–discussion 286–90. doi:10.1111/j.1365-2605.2005.00575.x

Herting, M.M., Gautam, P., Spielberg, J.M., Dahl, R.E., Sowell, E.R., 2015. A longitudinal study: changes in cortical thickness and surface area during pubertal maturation. PLoS ONE 10, e0119774. doi:10.1371/journal.pone.0119774

Hodes, G.E., Epperson, C.N., 2019. Sex Differences in Vulnerability and Resilience to Stress Across the Life Span. Biol Psychiat 86, 421–432. doi:10.1016/j.biopsych.2019.04.028

Howell, B.R., Godfrey, J., Gutman, D.A., Michopoulos, V., Zhang, X., Nair, G., Hu, X., Wilson, M.E., Sanchez, M.M., 2014. Social subordination stress and serotonin transporter polymorphisms: associations with brain white matter tract integrity and behavior in juvenile female macaques. Cereb Cortex 24, 3334–3349. doi:10.1093/cercor/bht187

Höistad, M., Barbas, H., 2008. Sequence of information processing for emotions through pathways linking temporal and insular cortices with the amygdala. NeuroImage 40, 1016–1033. doi:10.1016/j.neuroimage.2007.12.043

Koss, W.A., Lloyd, M.M., Sadowski, R.N., Wise, L.M., Juraska, J.M., 2015. Gonadectomy before puberty increases the number of neurons and glia in the medial prefrontal cortex of female, but not male, rats. Dev Psychobiol 57, 305–312. doi:10.1002/dev.21290

Lewis, G., Ioannidis, K., van Harmelen, A.-L., Neufeld, S., Stochl, J., Lewis, G., Jones, P.B., Goodyer, I., 2018. The association between pubertal status and depressive symptoms and diagnoses in adolescent females: A population-based cohort study. PLoS ONE 13, e0198804. doi:10.1371/journal.pone.0198804

Markham, J.A., Morris, J.R., Juraska, J.M., 2007. Neuron number decreases in the rat ventral, but not dorsal, medial prefrontal cortex between adolescence and adulthood. Neuroscience 144, 961–968. doi:10.1016/j.neuroscience.2006.10.015

McCormick, C.M., Green, M.R., 2013. From the stressed adolescent to the anxious and depressed adult: investigations in rodent models. Neuroscience 249, 242–257. doi:10.1016/j.neuroscience.2012.08.063

McEwen, B.S., Morrison, J.H., 2013. The brain on stress: vulnerability and plasticity of the prefrontal cortex over the life course. Neuron 79, 16–29. doi:10.1016/j.neuron.2013.06.028

McEwen, B.S., Nasca, C., Gray, J.D., 2016. Stress Effects on Neuronal Structure: Hippocampus, Amygdala, and Prefrontal Cortex. Neuropsychopharmacology 41, 3–23. doi:10.1038/npp.2015.171

Michopoulos, V., Higgins, M., Toufexis, D., Wilson, M.E., 2012. Social subordination produces distinct stress-related phenotypes in female rhesus monkeys. Psychoneuroendocrinology 37, 1071–1085. doi:10.1016/j.psyneuen.2011.12.004

Morgan, M.A., Schulkin, J., Pfaff, D.W., 2004. Estrogens and non-reproductive behaviors related to activity and fear. Neurosci Biobehav Rev 28, 55–63. doi:10.1016/j.neubiorev.2003.11.017

Naninck, E.F.G., Lucassen, P.J., Bakker, J., 2011. Sex differences in adolescent depression: do sex hormones determine vulnerability? J Neuroendocrinol 23, 383–392. doi:10.1111/j.1365-2826.2011.02125.x

Piekarski, D.J., Boivin, J.R., Wilbrecht, L., 2017. Ovarian Hormones Organize the Maturation of Inhibitory Neurotransmission in the Frontal Cortex at Puberty Onset in Female Mice. Curr Biol 27, 1735–1745.e3. doi:10.1016/j.cub.2017.05.027

Reding, K.M., Grayson, D.S., Miranda-Dominguez, O., Ray, S., Wilson, M.E., Toufexis, D., Fair, D.A., Sanchez, M.M., 2019. Effects of social subordination and estradiol on resting-state amygdala functional connectivity in adult female rhesus monkeys. J Neuroendocrinol e12822. doi:10.1111/jne.12822

Rincón-Cortés, M., Herman, J.P., Lupien, S., Maguire, J., Shansky, R.M., 2019. Stress: Influence of sex, reproductive status and gender. Neurobiol Stress 10, 100155. doi:10.1016/j.ynstr.2019.100155

Sallet, J., Mars, R.B., Noonan, M.P., Andersson, J.L., O’Reilly, J.X., Jbabdi, S., Croxson, P.L., Jenkinson, M., Miller, K.L., Rushworth, M.F.S., 2011. Social Network Size Affects Neural Circuits in Macaques. Science 334, 697–700. doi:10.1126/science.1210027

Sapolsky, R.M., 2005. The influence of social hierarchy on primate health. Science 308, 648–652. doi:10.1126/science.1106477

Schulz, K.M., Sisk, C.L., 2016. The organizing actions of adolescent gonadal steroid hormones on brain and behavioral development. Neurosci Biobehav Rev 70, 148–158. doi:10.1016/j.neubiorev.2016.07.036

Shansky, R.M., Rubinow, K., Brennan, A., Arnsten, A.F.T., 2006. The effects of sex and hormonal status on restraint-stress-induced working memory impairment. Behav Brain Funct 2, 8. doi:10.1186/1744-9081-2-8

Shansky, R.M., Woolley, C.S., 2016. Considering Sex as a Biological Variable Will Be Valuable for Neuroscience Research. J Neurosci 36, 11817–11822. doi:10.1523/JNEUROSCI.1390-16.2016

Skovlund, C.W., Mørch, L.S., Kessing, L.V., Lange, T., Lidegaard, Ø., 2018. Association of Hormonal Contraception With Suicide Attempts and Suicides. Am J Psychiatry 175, 336–342. doi:10.1176/appi.ajp.2017.17060616

Skovlund, C.W., Mørch, L.S., Kessing, L.V., Lidegaard, Ø., 2016. Association of Hormonal Contraception With Depression. JAMA Psychiatry 73, 1154–1162. doi:10.1001/jamapsychiatry.2016.2387

Snyder-Mackler, N., Sanz, J., Kohn, J.N., Brinkworth, J.F., Morrow, S., Shaver, A.O., Grenier, J.-C., Pique-Regi, R., Johnson, Z.P., Wilson, M.E., Barreiro, L.B., Tung, J., 2016. Social status alters immune regulation and response to infection in macaques. Science 354, 1041 – 1045. doi:10.1126/science.aah3580

Thapar, A., Collishaw, S., Pine, D.S., Thapar, A.K., 2012. Depression in adolescence. Lancet 379, 1056–1067. doi:10.1016/S0140-6736(11)60871-4

Tottenham, N., Galvan, A., 2016. Stress and the adolescent brain: Amygdala-prefrontal cortex circuitry and ventral striatum as developmental targets. Neurosci Biobehav Rev 70, 217–227. doi:10.1016/j.neubiorev.2016.07.030

Twenge, J.M., Cooper, A.B., Joiner, T.E., Duffy, M.E., Binau, S.G., 2019. Age, period, and cohort trends in mood disorder indicators and suicide-related outcomes in a nationally representative dataset, 2005-2017. J Abnorm Psychol 128, 185–199. doi:10.1037/abn0000410

van Duijvenvoorde, A.C.K., Westhoff, B., de Vos, F., Wierenga, L.M., Crone, E.A., 2019. A three-wave longitudinal study of subcortical-cortical resting-state connectivity in adolescence: Testing age-and puberty-related changes. Hum Brain Mapp 40, 3769–3783. doi:10.1002/hbm.24630

Weintraub, A., Singaravelu, J., Bhatnagar, S., 2010. Enduring and sex-specific effects of adolescent social isolation in rats on adult stress reactivity. Brain Res 1343, 83–92. doi:10.1016/j.brainres.2010.04.068

Wierenga, L.M., Bos, M.G.N., Schreuders, E., vander Kamp, F., Peper, J.S., Tamnes, C.K., Crone, E.A., 2018. Unraveling age, puberty and testosterone effects on subcortical brain development across adolescence. Psychoneuroendocrinology 91, 105–114. doi:10.1016/j.psyneuen.2018.02.034

Wilson, M.E., 2016. An Introduction to the Female Macaque Model of Social Subordination Stress, in: Social Inequalities in Health in Nonhuman Primates, Developments in Primatology: Progress and Prospects. Springer, Cham, Cham, pp. 9–24. doi:10.1007/978-3-319-30872-2_2

Wilson, M.E., Bounar, S., Godfrey, J., Michopoulos, V., Higgins, M., Sanchez, M., 2013. Social and emotional predictors of the tempo of puberty in female rhesus monkeys. Psychoneuroendocrinology 38, 67–83. doi:10.1016/j.psyneuen.2012.04.021

Wilson, M.E., Chikazawa, K., Fisher, J., Mook, D., Gould, K.G., 2004. Reduced growth hormone secretion prolongs puberty but does not delay the developmental increase in luteinizing hormone in the absence of gonadal negative feedback. Biol Reprod 71, 588–597. doi:10.1095/biolreprod.104.027656

